# RPSLearner: A novel approach based on random projection and deep stacking learning for categorizing NSCLC

**DOI:** 10.1101/2025.05.01.651699

**Authors:** Xinchao Wu, Jieqiong Wang, Shibiao Wan

**Affiliations:** Department of Genetics, Cell Biology and Anatomy, University of Nebraska Medical Center, Omaha, NE; Department of Neurological Sciences, University of Nebraska Medical Center, Omaha, NE

**Keywords:** Lung cancer subtype prediction, Random projection, Stacking Learning, Machine learning, Transcriptomics

## Abstract

**Background:** Lung cancer is the leading cause of cancer death, and non-small cell lung cancer (NSCLC) comprises the largest subtype with most cases. Lung adenocarcinoma (LUAD) and lung squamous cell carcinoma (LUSC) are two NSCLC subtypes that pose challenges for accurate diagnosis using conventional methods. Existing methods are histological examination and imaging which lacks definitive histologic features and requires intense time.

**Methods:** To address these concerns, we propose RPSLearner, which combines Random Projection (RP) for dimensionality reduction and stacking ensemble learning to accurately predict lung cancer subtypes. Specifically, multiple independent RP matrices were first generated to project the high-dimensional RNA-seq data into lower-dimensional space, whose features were subsequently concatenated. After that, we fed the fused features into a stack of diverse base classifiers and integrated the predictions from base models via a deep linear layer network.

**Results:** Benchmarking tests on 1,333 NSCLC patients demonstrated that RPSLearner outperformed state-of-the-art approaches for lung cancer subtype classification. Specifically, RPSLearner efficiently preserved sample-to-sample distances even after significant dimension reduction, and the meta-model in RPSLearner yielded consistently higher accuracy, F1 and AUC scores than individual base models and state-of-the-art approaches for lung cancer subtyping.

Besides, the feature fusion method applied in RPSLearner shown better performance than conventional scores ensemble methods.

**Conclusion:** We developed a novel stacking learning method called RPSLearner which combines RP and stacking learning, enabling efficient and accurate identification of NSCLC subtypes. RPSLearner is a promising lung cancer subtyping model for downstream lung cancer clinical diagnosis and personalized treatment, and the framework holds the potentiality to be extended to subtyping of other types of cancer.

## Introduction

Lung cancer remains one of the leading causes of cancer-related mortality worldwide, accounting for a substantial proportion of cancer diagnoses and deaths each year^1,2^. Non-small cell lung cancer (NSCLC) constitutes approximately 85% of all lung cancer cases, with subtypes such as lung adenocarcinoma (LUAD) and lung squamous cell carcinoma (LUSC), exhibiting distinct molecular and clinical characteristics^3^. Accurate identification of NSCLC subtypes is critical for prognosis determination, therapeutic decision-making, and the development of personalized treatment strategies^4^. Traditional diagnostic methods, including histopathological examination and imaging techniques^5^, while valuable, often lack the precision required for nuanced subtype differentiation, particularly in early-stage disease^6^. Besides, histological examination increases the risk of lung cancer metastasis due to physical damage^7^.

In recent years, advancements in high-throughput genomic and transcriptomic technologies have revolutionized our understanding of the molecular mechanisms of lung cancer, enabling the identification of specific biomarkers and genetic signatures associated with different NSCLC subtypes^8^. Among these techniques, transcriptomic profiling provides a dynamic and quantitative snapshot of gene expression, thereby serving as a more direct indicator of the cellular functional state^9^. However, the analysis of such high-dimensional omics data presents significant computational and methodological challenges. The complexity and volume of transcriptomic profiles require robust feature selection and dimensionality reduction techniques to extract meaning patterns while mitigating the risk of overfitting and enhancing model generalizability^10^.

Machine learning (ML) approaches have emerged as powerful tools for integrating and analyzing large-scale biological data, offering the potential to improve the accuracy and efficiency of lung cancer subtype prediction. For instance, Huang et al.^11^ utilized logistic regression (LR) with LASSO shrinkage for predicting NSCLC subtypes based on RNA-seq data. Their model demonstrated superior performance compared to LR using principal components and K-nearest neighbor (KNN) algorithms. Similarly, Yuan et al.^12^ applied a method called Monte Carlo feature selection (MCFS) in combination with support vector machine (SVM) for subtype stratification based on microarray gene expression data, showing better results than the random forest (RF) classifier. Although these studies illustrated the effectiveness of machine learning in NSCLC subtype classification, significant challenges remain. In particular the high dimensionality of omics data can lead to overfitting and reduced generalizability, and inter-sample variability further complicates the creation of robust predictive models^13^. Recently we have witnessed the significant development of multiple deep learning approaches^14,15^ tailored to cancer subtype diagnosis combining histological images and genomics data, which have demonstrated promising results for cancer subtyping. However, these types of methods require the availability of transcriptomics data paired with histological staining imaging data, which are very costly or laborious to obtain. In this case, we focus on developing a novel approach solely based on transcriptomics data for cost-effective and accurate NSCLC subtype classification.

To address these challenges, we developed RPSLearner combining Random Projection (RP) for dimensionality reduction and stacking ensemble learning to accurately predict lung cancer subtypes. In detail, by leveraging multiple independent RP matrices and fusing these reduced features, RPSLearner can capture essential sample-to-sample distances^16,17^. Subsequently, RPSLearner uses stacking learning framework to integrate diverse heterogeneous base classifiers with a deep linear network as a meta-model^18,19^. The stacking framework not only consolidates complementary predictive insights but also yields improved performance metrics over conventional methods. Ultimately, in this study, we demonstrated that RPSLearner significantly enhance NSCLC subtype classification and offered a versatile framework that may be adapted for other cancer subtyping applications.

## Results

### The design of RPSLearner

To accurately identify subtypes of NSCLC, we presented RPSLearner, a novel computational framework that integrates RP with stacking ensemble learning. The whole process of the method was shown in **Fig. 1a**. RPSLearner leverages RP’s capabilities of dimensionality reduction to deal with the high-dimensional nature of transcriptomic data^20^. Specifically, it applied multiple random projections to patients’ transcriptomic profiles to extract meaningful hidden information, significantly decreasing the feature space while preserving the intrinsic distances between samples. This approach facilitated the handling of large-scale omics datasets, ensuring computational efficiency without compromising the integrity of the underlying biological signals.

**Figure 1.**
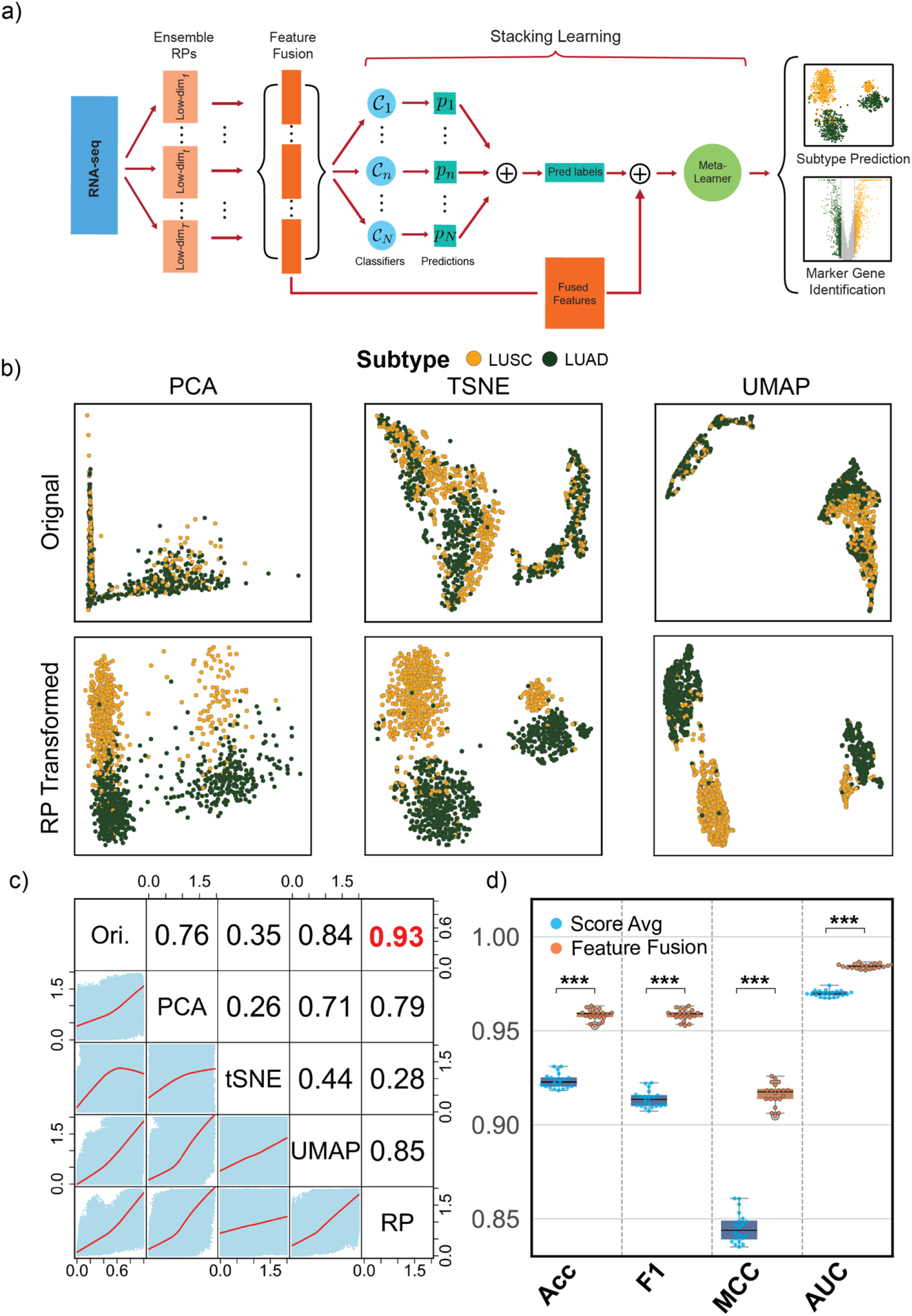
The overview of lung cancer TCGA datasets and the RPSLearner algorithm. (**a**) The computational pipeline of the RPSLearner. (**b**) Low-dimension visualization of original data and RP-transformed data through diverse techniques. (**c**) The correlative analysis across diverse dimensionality reduction methods. (**d**) The performance of score average strategy against feature fusion. Ori.: Original, PCA: Principle Component Analysis, tSNE: t-distributed Stochastic Neighbor Embedding, UMAP: Uniform Manifold Approximation and Projection. RP: Random Projection. Statistical significance level denoted as: ns (not significant) or P ≥ 0.05, * for P < 0.05, ** for P < 0.01, *** for P < 0.001.

### RPSLearner efficiently preserved sample-to-sample distances

To demonstrate the effectiveness of RP, we applied PCA^21^, t-SNE^22^, and UMAP^23^ to visualize the RP-transformed NSCLC RNA-seq data and compared them against original data (**Fig. 1b**). Across diverse techniques, the RP-transformation, fusing multiple RP dimension reduction results, exhibited clear subtype identities with distinct cluster groups, suggesting that effectively extract meaningful features in lung cancer subtype prediction and simplify the prediction task.

To further validate the performance of RP for dimensionality reduction, we next evaluated RP against PCA, t-SNE, and UMAP for NSCLC RNA-seq data. **Fig. 2b** compared Pearson correlation coefficients and distance-fitting curve between original and projected feature vectors.

**Figure 2.**
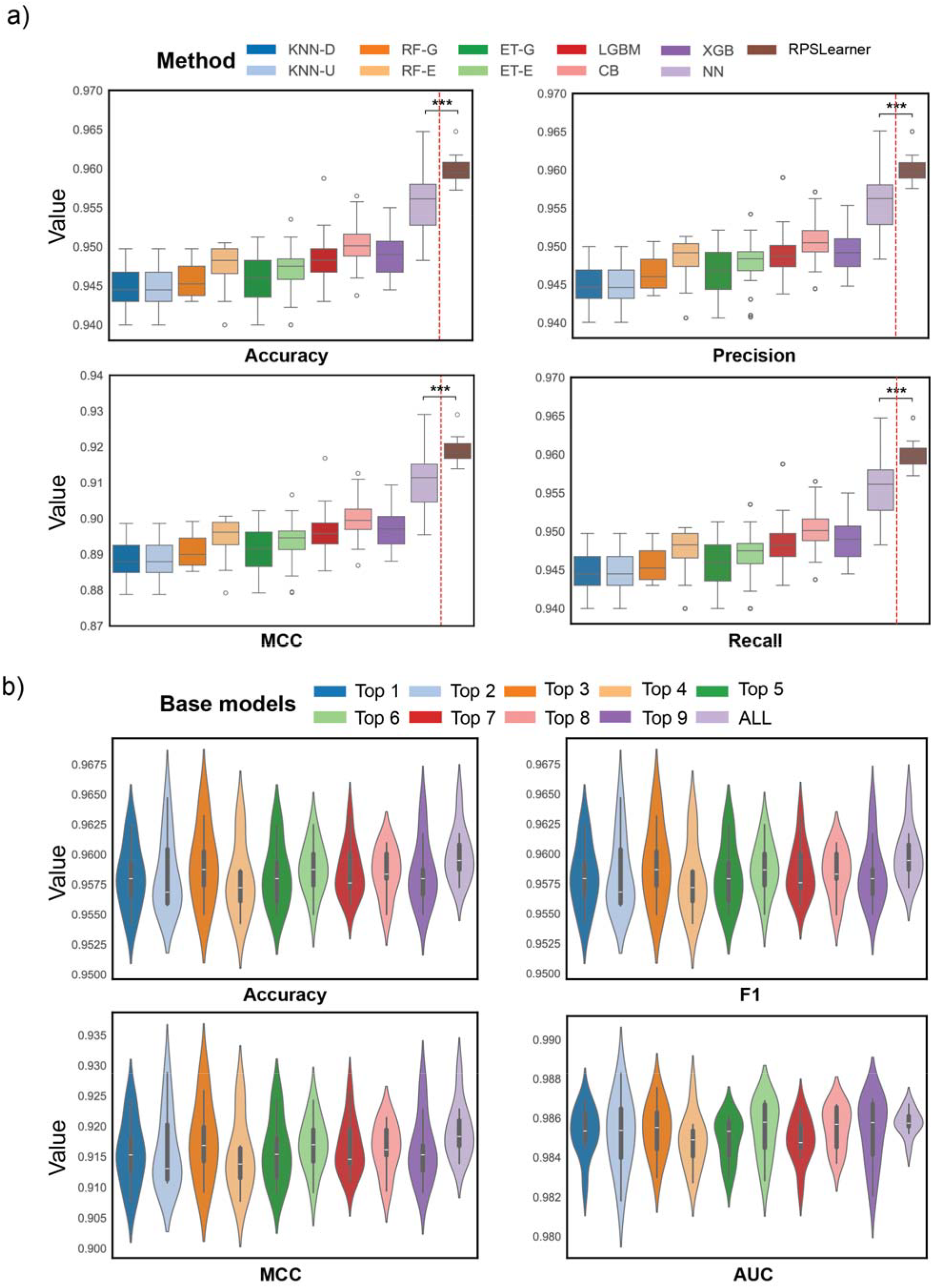
The performance of RPSLearner in lung cancer subtype prediction. (**a**) The comparison of performance of diverse individual base model and stacked model, RPSLearner. (**b**) The comparison of performance of various combination of base models in stacking learning, top k selection was based on results of previous individual base model comparison, ALL represented using all ten models. KNN-D: KNN in distance metrics, KNN-U: KNN in uniform metrics, RF-G: RF in Gini metrics, RF-E: RF in entropy metrics, ET-G: ExtraTree in Gini metrics, ET-E ExtraTree in entropy metrics, LGBM: LightGBM, CB: CatBoost, XGB: XGBoost, NN: neural network. Statistical significance level denoted as: ns (not significant) or P ≥ 0.05, * for P < 0.05, ** for P < 0.01, *** for P < 0.001.

RP consistently maintained higher correlations across various dimensions up to 0.94 (**Supplementary Fig. 2**, more effectively preserving sample-to-sample relationships. UMAP showed the second-best correlation (0.84) but lagged RP. PCA and t-SNE were less effective in preserving pairwise distances for this dataset. These findings underscore RP’s suitability for retaining meaningful data patterns in a reduced-dimension space.

To demonstrate the effectiveness of the stacking learning method in RPSLearner, we compared two ways of leveraging multiply RPs, feature fusion and score ensemble. Conventionally, methods applied score ensemble, which averaging the prediction probabilities from independent runs as the final score. Feature fusion concatenates RP-transformed features for each randomly generated feature and feed them into stacking learning pipeline. We compared the performance between modified RanBALL^16^, which originally used for leukemia subtypes prediction applied ensemble strategy, and RPSLearner, using feature fusion. Through 20 times 5-fold CV, we demonstrated feature fusion yields higher accuracy, F1 score, MCC and AUC than score ensemble (**Fig. 1d**), suggesting that integrating projected features directly exploits complementary representations more efficiently than simple probability averaging.

### Stacking meta learning outperforms individual base learning models

We next tested RPSLearner using different projected dimension sizes (from 100 to 1,400) to find optimal performance (**Supplement Fig 3a**). With 10 times 5-fold cross validation (CV), it demonstrated that the accuracy and other metrics, improved from 100 to 400 dimensions and then achieved stabilized. Subsequently, we explored to use multiple independent RP matrices (**Supplement Fig. 3b**). With the same setting using 10 times 5-fold CV, it showed that accuracy and other metrics steadily rose as the number of RPs used up to around 25, reaching 0.961 in accuracy, 0.96 in F1, 0.92 in MCC, and 0.986 in AUC on average. Beyond 25 RPs, the performance gains were marginal, suggesting that moderate RP ensembles sufficiently capture complex signals.

To evaluate the performance of stacking learning, we compared the performance among each individual base model and NN-based meta-model. We assessed them using 10 times 5-fold CV with 20 individual 400-dimensional RP matrices. **Fig. 2a** showed meta-model achieved the best performance across all metrics. The NN base classifier performed the second best, but stacking meta-model delivered a consistent and significant boost in accuracy, F1, and MCC. For AUC, individual strong learners attained high scores around 0.985, leaving less room for stacking gains to show significant improvement. Nevertheless, the stacking approach never degraded performance.

To deeper examine the effectiveness of stacking learning, we compared the performance across various composite of base models in used. We assessed with the same RP setting as above, and **Fig 2b** exhibited that the performance of using all base models obtained smallest variance while was no less than the best performance of base model combination using top 3. In other words, it enhanced the robustness of the model prediction.

### RPSLearner outperformed state-of-the-art approaches for lung cancer subtype identification

To demonstrate that RPSLearner could outperform state-of-the-art methods, we benchmarked RPSLearner against previously published methods for NSCLC subtype prediction (**Fig. 3a**). Approaches includes MCFS + support vector machine (SVM), MCFS + RF, Lasso + SVM, Lasso + RF, ANOVA + SVM^24^, ANOVA + RF and a Lasso-based developed by Huang et al.^11^ obtained accuracy ranging from around 0.93 to 0.95. By contrast, RPSLearner achieved around 0.96 accuracy, which is significantly higher than SOTA method. The consistent and significant improvement also shown in other metrics. Besides, we further identify differential expression genes (DEGs) based on RPSLearner prediction (**Fig. 3b, d**) There are many well-established biomarkers of LUAD and LUSC included in the identified DEGs. For example, KRT5, KRT6A, KRT14, and DSG3 in LUSC^25^, and NKX2.1^26^, SFTA2^27^ in LUAD have been evidently proved directly correlated to the lung cancer diagnosis and progression. Additionally, we compared the DEGs identified with prediction of other methods and ground-truth (**Fig. 3c**). Through RPSLearner prediction, there is a larger overlap between RPSLearner with ground-truth, and RPSLearner could find more DEGs than ground-truth. These findings highlight the accuracy of the prediction, as well as potential novel molecular signatures identified through RPSLearner prediction.

**Figure 3.**
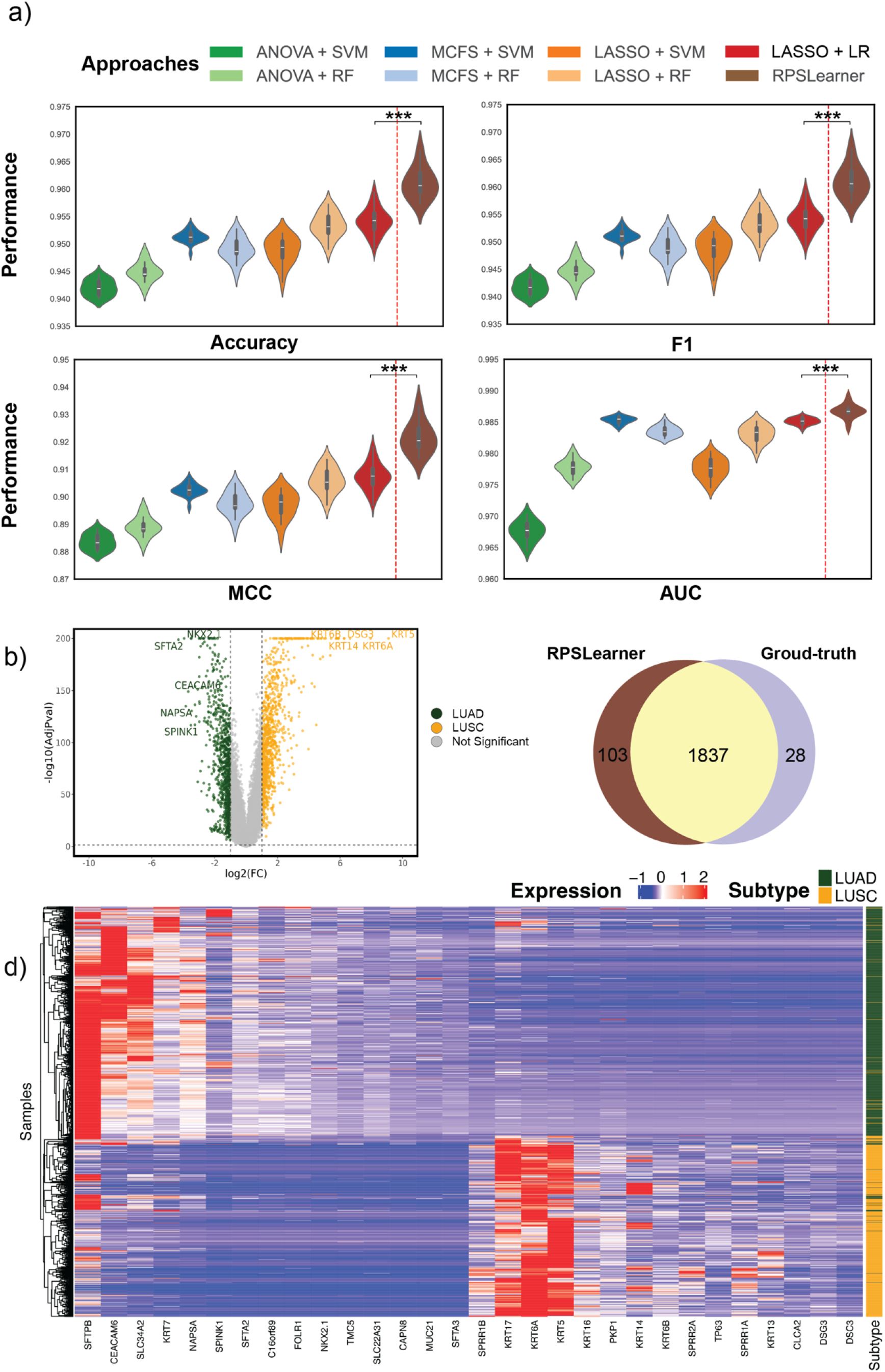
RPSLearner outperforms state-of-the-art methods with interpretable results. **(a**) The benchmark of methods for lung cancer subtype prediction. (**b**) The volcano plot illustrates the significance of the DE genes of each subtype. (**c**) Comparison among differential expression genes (DEGs) identified based on RPSLearner and ground-truth. (**d**) Expression of the DEGs extracted from RPSLearner prediction. Statistical significance level denoted as: ns (not significant) or P ≥ 0.05, * for P < 0.05, ** for P < 0.01, *** for P < 0.001. MCFS: Monte Carlo Feature Selection.

## Discussion

In this study, we introduced RPSLearner, a novel approach that combines RP for dimensionality reduction and stacking learning framework to aggregate predictions from diverse base models. RPSLearner utilized RP to reduce the dimensionality of transcriptomics data. Unlike other dimensionality reduction approaches that distorted intrinsic sample-to-sample relationships, RP applied in RPSLearner was able to capture meaningful and accurate latent features from the transcriptomics data, efficiently reduce dimensionality while preserve intrinsic distances among samples. This was crucial for maintaining the structural integrity of the data, which is essential for accurate subtype prediction.

RPSLearner applied feature fusion which concatenated multiple independent RP reduced features for a richer representation learning. Compared to score ensemble strategies used in conventional RP-based algorithms that merely integrated the model scores from multiple RP reduced results, feature fusion could represent a richer aspect of hidden information since it integrated multi-view latent features, and each reduced feature was independently generated from the original transcriptomics data. These results suggested that it could efficiently exploit variances from multiple RP matrices, achieving a more representative latent features for lung cancer subtype prediction. These results demonstrated exploiting multi-scale feature space could enhance the effectiveness and generalization ability of the model.

Another unique advantage of RPSLeaner is the stacking framework. RPSLearner leverages a meta-learner, specifically a deep linear neural network to learn the fused features combined multiple RP reduced results and diverse conventional machine learning models. The stacking framework leveraged the complementary strengths of diverse base models as well as the insights from the meta-learner. The meta-leaner could learn the utilize the knowledge from other leading to improved performance across multiple metrics. Our results emphasized that the predictions of meta-model are better than that from each individual base model. The use of stacking learning framework improves the robustness and capacity of the prediction model through leveraging various base models with heterogeneous ability to explore the same data.

RPSLearner demonstrated superior performance compared to state-of-the-art lung cancer subtype prediction methods. We tested the other methods, such as MCFS + SVM from Huang et al.^12^ which applied MCFS for feature selection, and SVM for cancer subtype prediction. Comparative analysis revealed that RPSLearner consistently showed higher performance across various metrics. A reasonable explanation to RPSLearner’s superior performance was that the application of fused features and stacking learning to leverage both RP transformed features and predictions from diverse base models rather than simply averaging the prediction results.

RPSLearner exhibited biological interpretable to identify characteristic biomarkers from NSCLC subtypes. RPSLearner not only extract meaningful hidden features from transcriptomics data but also offers insights about the detection of novel potential biomarkers. These results highlighted its potential for clinical application. Besides, the method could be extended to more cancer types that each cancer type may present distinct molecular and genomic characteristics, making is essential to evaluate the effectiveness in universal cancer subtypes prediction^28^. Moreover, the method could also be extended to other tasks related to the cancer diagnosis such as cancer progression prediction, where the goal shifts from classifying subtypes to predict tumor growth or patient prognosis over time^29^. By incorporating longitudinal data, such as multiple time points of gene expression data, RPSLearner could provide valuable insights into disease progression, ultimately informing personalized treatment strategies^30^.

In addition, we also note that there are potential future research directions that can be explored to improve the performance of RPSLearner. First, we need to expand the dataset usage to validate the efficacy of the model, which is necessary for translating these methods into clinical practice. This plan remains challenging due to the high dimensionality of data and the need for models that are both accurate and interpretable^31^, and considering the high computational cost of stacking framework^32^. It requires us to develop more efficient approaches. To solve this concern, we need to fully utilize high-performance computational resources such as GPU to accelerate the computation processing.

## Conclusion

In this paper, to address the concerns in NSCLC subtype prediction, we developed RPSLearner which combines RP and stacking learning for effective and accurate classification. It effectively reduced the dimensionality while preserving sample-to-sample distances through RP and integrated fused features and predictions from diverse models through stacking learning. RPSLearner succeeds in boosting classification prediction with higher accuracy, F1 and AUC metrics than conventional machine learning models and state-of-the-art methods. RPSLearner utilized feature fusion strategy which exhibited better performance than score ensemble approaches in subtype prediction. RPSLearner’s results are interpretable that the expression of DEGs aligns well with the published literature, which also offering insights about potential novel biomarkers. This framework could be potentially extended to subtype identification of other cancers.

## Methods

### Dataset

This study collected RNA-seq data for LUAD and LUSC retrieved from the NCI Genomic Data Commons (GDC) database^33^ with 9 different data sources (**Supplement Fig. 1**). We excluded samples with incomplete expression profiles and yielded a final dataset of 1,333 cases, with 731 LUAD and 602 LUSC cases, respectively. The raw read counts of all RNA-seq samples were normalized to Transcripts Per Million (TPM).

### RPSLearner

Specifically, considering a data matrix **X** of size *n* × *d*, where *n* was the number of samples and *d* was the original dimensionality. RP found a random matrix **R** ∈ ℝ*^d×k^*, with *k* « *d*, such that the transformed data matrix***Y*** = ***X*** ·**R**, ∈ ℝ *^n ×k^*, captured the hidden geometry of the original data **X** in the reduced dimension *k*.

One common choice of **R** is a Gaussian RP matrix^34^. Each entry *r*_ij_ in **R** is independently sampled from a Gaussian distribution 𝒩(0, *σ*^2^), where *σ*^2^ is set to 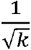, ensuring that the resulting projected vectors are suitable to Johnson-Lindenstrauss Lemma. In other words,

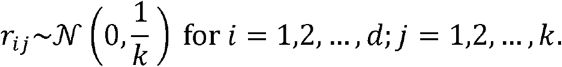

Then for a data point***x*** ∈ ℝ *^d^*, its - dimensional projection is ***y*** = ***x*** ·**R**. With high probability, the Euclidean distances among samples in the projected space ℝ *^k^* will be close to those in original space ℝ *^d^*.

Following RP-based dimensionality reduction, the transformed feature vectors were fed into diverse machine-learning base models, each generating unique predictive labels. These base models encompassed a range of algorithms, including K-Nearest Neighbors (KNN)^35^, Random Forest (RF)^36^, LightGBM^37^, Extra Trees^38^, Catboost^39^, XGBoost^40^, Neural Networks^41^, thereby capturing a wide spectrum of data patterns and interactions. Then, RPSLearner aggregated the predictions from these base models with transformed feature vectors using a stacking meta-learner, specifically a multi-layer perceptron (MLP), which synthesized the individual predictions and features to enhance overall classification performance. This stacking strategy not only mitigated the biases inherent in individual models but also leveraged their complementary strengths, resulting in a more robust and accurate subtype prediction.

In our stacking framework, we employed a diverse set of base classifiers, each generating its own prediction. Let ***x*** ∈ ℝ*^d^* denote an input feature vector from the dataset and let the set of possible class labels be *ℓ* = {1,2,…*C*}. Each base classifier *f_i_* (·) returns its predicted label 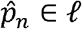.

The stacking method required a meta learner model to integrate the prediction from various base models. Let there be *B* base models {*f*_1_,*f*_1_,…*f_N_*}. Each model *f_N_* could output a predicted label 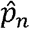.. For numerical stability and richer information, we concatenated of all base model predictions, appending the original input ***x*** to form a composite feature 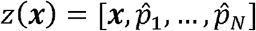. The final prediction *ŷ* could be formulated as:

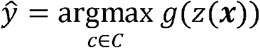

Where *g*(*z*(***x***)) represented the stacking meta-learning model and argmax function aims to get the prediction label with highest probability.

## Authors’ Contributions

X.W.: data collecting, data preprocessing, model development, data analysis and interpretation, manuscript preparation, editing, and review. J.W.: manuscript editing and review. S.W.: Study concept and design, manuscript editing and review.

## Data availability

The RNA-seq data of lung cancer samples can be publicly accessed from the NCI GDC database: https://portal.gdc.cancer.gov.

## Code availability

The RPSLearner Python package can be accessed at https://github.com/wan-mlab/RPSLearner.

